# Investigations of human myosin VI targeting using optogenetically-controlled cargo loading

**DOI:** 10.1101/068965

**Authors:** Alexander R. French, Tobin R. Sosnick, Ronald S. Rock

## Abstract

Myosins play countless critical roles in the cell, each requiring it to be activated at a specific location and time. To control myosin VI with this specificity, we created a novel optogenetic tool for activating myosin VI by fusing the light-sensitive *Avena sativa* phototropin1 LOV2 domain to a peptide from Dab2 (LOVDab), a myosin VI cargo protein. Our approach harnesses the native targeting and activation mechanism of myosin VI, allowing direct inferences on myosin VI function. LOVDab robustly recruits human full length myosin VI to various organelles *in vivo* and hinders peroxisome motion in a light-controllable manner. LOVDab also activates myosin VI in an *in vitro* gliding filament assay. Our data suggest that protein and lipid cargoes cooperate to activate myosin VI, allowing myosin VI to integrate Ca^2+^, lipid, and protein cargo signals in the cell to deploy in a site-specific manner.

**Significance Statement:** Myosins are a broad class of motor proteins that generate force on actin filaments and fulfill contractile, transport, and anchoring roles. Myosin VI, the only myosin to walk toward the pointed end of actin filaments, is implicated in cancer metastasis and deafness. Intriguingly, myosin VI may play both transport and anchoring roles, depending upon where it is activated in the cell. Here we develop an optogenetic tool for studying myosin VI activation with high spatial and temporal resolution. Our approach photoactivates unmodified myosin VI through its native cargo pathway, enabling investigation of motor function and activation partners with minimal perturbation. This approach allows us to detect how and where myosin VI integrates multiple protein and second messenger signals to activate.

## Introduction

Motor proteins play countless roles in biology, each requiring the motor to be recruited and activated at a particular time and place inside the cell. To dissect these multiple roles, we must develop tools that allow us to control the recruitment and activation process. One promising technique for achieving this goal is through optogenetics (1). Optogenetics involves the engineering and application of <u>opt</u>ically-controlled, <u>genetic</u>ally encoded proteins, and is transforming the fields of neuro- and cell biology (2, 3). A major benefit of optogenetics is that proteins are activated using light, which allows for high temporal and spatial control over a protein of interest.

Myosin VI is a motor protein whose study could particularly benefit from optogenetic control. It is the only myosin known to walk toward the pointed end of actin filaments (4, 5). This property enables it to perform a diverse array of cellular functions including cell division, endosome trafficking, autophagy, and Golgi and plasma membrane anchoring (6–9). Myosin VI is also auto-inhibited, a property that is commonly found in other myosins (10). When myosin VI binds to cargo through specialized adaptor proteins, this auto-inhibition is relieved through a poorly understood mechanism likely involving the disruption of an interaction between its cargo binding domain (CBD) and the myosin head (11, 12). Dissociation of the CBD from the head both frees the head to bind tightly to actin and exposes dimerization sites throughout the tail domain of myosin VI, allowing it to become a processive dimer (13–15). In some cases, myosin VI could conceivably function as a monomer, for example when fulfilling its role as a membrane tether during spermatid individualization (16). If this is the case, more work is needed to elucidate the cellular signals that determine its oligomeric state at each site of action.

Myosin VI has two classes of cargo proteins that bind to distinct, conserved motifs on the myosin VI C-terminus (17). Disabled2, or Dab2, belongs to the class of cargo proteins that bind to a conserved WWY site on myosin (18). Optineurin (OPTN) is a member of the second class that binds to a conserved RRL motif (19). Binding to members of either of these classes of cargo protein is thought to relieve the auto-inhibition through the mechanism described above. Recently, however, myosin VI’s activation mechanism has been found to be more elaborate, with evidence now showing that myosin VI binds specifically to the signaling lipid PI(4,5)P_2_ as well as being activated using Ca^2+^ (18, 20). It remains unclear how these small molecule signals influence the cargo protein pathway above, or whether they represent a distinct pathway for myosin VI activation. These outstanding questions on its activation mechanism, together with its many roles throughout the cell, make myosin VI an ideal target for optogenetic study.

One highly successful optogenetic strategy is to use or modify naturally-occurring light sensitive proteins that directly alter a cellular property of interest, such as using channel rhodopsin to manipulate membrane ion gradients (1). Another method is to fuse a target protein to a naturally light-sensitive protein that undergoes a conformational change in response to light. The new fusion protein is designed such that it sterically blocks the target protein’s binding to its partner in the dark state, but releases it in the lit state.

This latter engineering strategy has been employed using a variety of light-sensitive proteins, including phytochromes (21, 22), cryptochromes (23), photoactive yellow protein (PYP) (24), and the Light, Oxygen, Voltage sensing domain 2, or LOV2, from *Avena sativa* phototropin1 (25–27). Each system has its own benefits (1), but we proceeded with the LOV2 domain due to its ability to cage a peptide in the dark, and expose it to binding effectors in the light. The ~150 AA domain binds a flavin mononucleotide (FMN) and undergoes a conformational change involving the unfolding of its N- and C-terminal helices upon absorbing blue light (λ_max_ ≈ 447nm) (28). Specifically, absorption leads to the formation of a covalent bond between the FMN’s C4a atom and the sulfur atom of the catalytic cysteine residue C450 (29, 30). This event triggers the unfolding of both the A′α helix at its N-terminus and the larger Jα helix at its C-terminus (31, 32). When the light stimulus is removed, the protein-FMN bond spontaneously hydrolyses in the dark on the seconds to minutes time scale followed by rapid refolding of the A’α and Jα helices, thus completing the photocycle (30–32). The degree of this conformational change and the rate at which the helices refold in the dark can be independently altered by mutation (32, 33) which can be useful for tuning a photoswitch (25, 26).

Previously, we engineered the LOV2 domain to control DNA binding (34) as well as to sterically block, or “cage,” a small PDZ domain-binding peptide that was fused to the C-terminal Jα helix (26). In the latter system, the light-induced unfolding of the Jα helix uncages the peptide and recruits PDZ domain-tethered proteins *in vivo*. This strategy has been utilized by others for the control of a variety of protein-protein interactions (35) including most recently epigenetic modifications (36).

Here we apply a similar uncaging strategy to engineer a LOV2 fusion protein that can recruit myosin VI in a light controllable manner. A short peptide region from the myosin VI cargo protein Dab2 (Dab2^pep^) has been crystallized in complex with the cargo binding domain (CBD) of myosin VI, and shown to recruit the myosin VI tail *in vivo* (15). Phichith *et al*. independently were able to activate full-length porcine myosin VI *in vitro* using high concentrations of a Dab2 truncation that included Dab2^pep^ (13). Additionally, ITC experiments suggest that the N-terminal α-helix of Dab2^pep^ contains most of its binding affinity for myosin VI, making Dab2^pep^ a suitable target for caging by the LOV2 domain.

Here we engineer and apply an optogenetic activator for full-length human myosin VI using a LOV2-Dab2^pep^ fusion protein termed “LOVDab”. LOVDab robustly recruits myosin VI to a variety of cellular organelles in a light-dependent manner. Recruitment of myosin VI to peroxisomes slows their velocity, demonstrating myosin VI activation upon recruitment *in vivo* that is consistent with an anchoring role for myosin VI. We also purify full-length human myosin VI and demonstrate its photoactivation *in vitro* using a modified gliding filament assay. Lastly, we interrogate the interplay of lipid and cargo proteins in activating myosin VI, showing for the first time that the presence of PI(4,5)P_2_ enhances myosin VI’s affinity for protein cargo *in vitro*. We also find that the amount of myosin VI recruited by LOVDab to secretory pathway membranes is independent of LOVDab concentration, suggesting that other factors must influence myosin VI’s affinity for LOVDab at each membrane. Our results support a new model of myosin VI activation where myosin VI integrates lipid, Ca^2+^, and protein cargo signals to activate in a site-specific manner.

## Results

### LOVDab design and optimization

To obtain light-controlled activation of myosin VI, we fused the portion of its cargo protein Dab2 (res. 674-711, Dab2^pep^) that binds to the myosin VI cargo binding domain (CBD) to the C-terminus of the LOV2 Jα helix. In the crystal structure of the CBD-Dab2^pep^ complex, two CBDs and two Dab2^pep^ molecules form a tetrameric structure (Fig. 1*A*). Each Dab2^pep^ in the tetramer contains two helical regions that contact the CBDs. The two helices bind to opposite CBDs so that each Dab2^pep^ binds across both CBDs in the tetramer. The modest interface between the two CBDs in the tetramer suggests that two Dab2^pep^s promote myosin VI dimerization by tethering two myosin VI molecules in close proximity to each other, disrupting the interaction of the CBD to the myosin VI head domain and promoting myosin VI dimerization at various sites along its tail domain (13–15). Whether Dab2^pep^ directly competes with the interaction between the CBD and the myosin VI head is unknown. ITC measurements indicate that the majority of the binding affinity for the CBD lies in the N-terminal helix of Dab2^pep^ (15). Taken together, these observations suggest that light-induced uncaging of the N-terminal region of two Dab2^pep^ peptides may be sufficient to control myosin VI dimerization. Furthermore, the 2:2 stoichiometry of the CBD_2_:Dab2^pep^_2_ tetramer results in a binding reaction dependent on the square of the concentration of Dab2^pep^, which should amplify the switching capability of our design.

Myosin VI cargo proteins tether myosin VI to membrane-bound organelles *in vivo*. Accordingly, we tested our designs using an assay that would recruit myosin VI to various membranes in the cell (Fig. 1*B*). Mitochondria were an ideal first choice for such an assay since proteins can be targeted to the mitochondrial membrane by fusing the Tom70 transmembrane helix (Tom70^helix^) to their N-termini (26). Additionally, myosin VI is not known to act on mitochondria, and overexpression of myosin VI in the cell should therefore not lead to increased background binding of myosin VI to this organelle. We thus cotransfected HeLa cells with Tom70^helix^-Fluorescent Protein (FP)-LOVDab and −LOVDab_+ctrl_ fusion proteins and full-length human myosin VI tagged with a complimentary FP on its N-terminus. Because the absorption λ_max_ of EGFP is similar to that of LOV2, we alternate which of two FPs, mCherry or EGFP, is on myosin VI and which is on LOVDab. The protein we intend to watch in both the light and the dark states of LOV2 is fused to mCherry in order to avoid simultaneous excitation of LOV2.

**Fig. 1.**
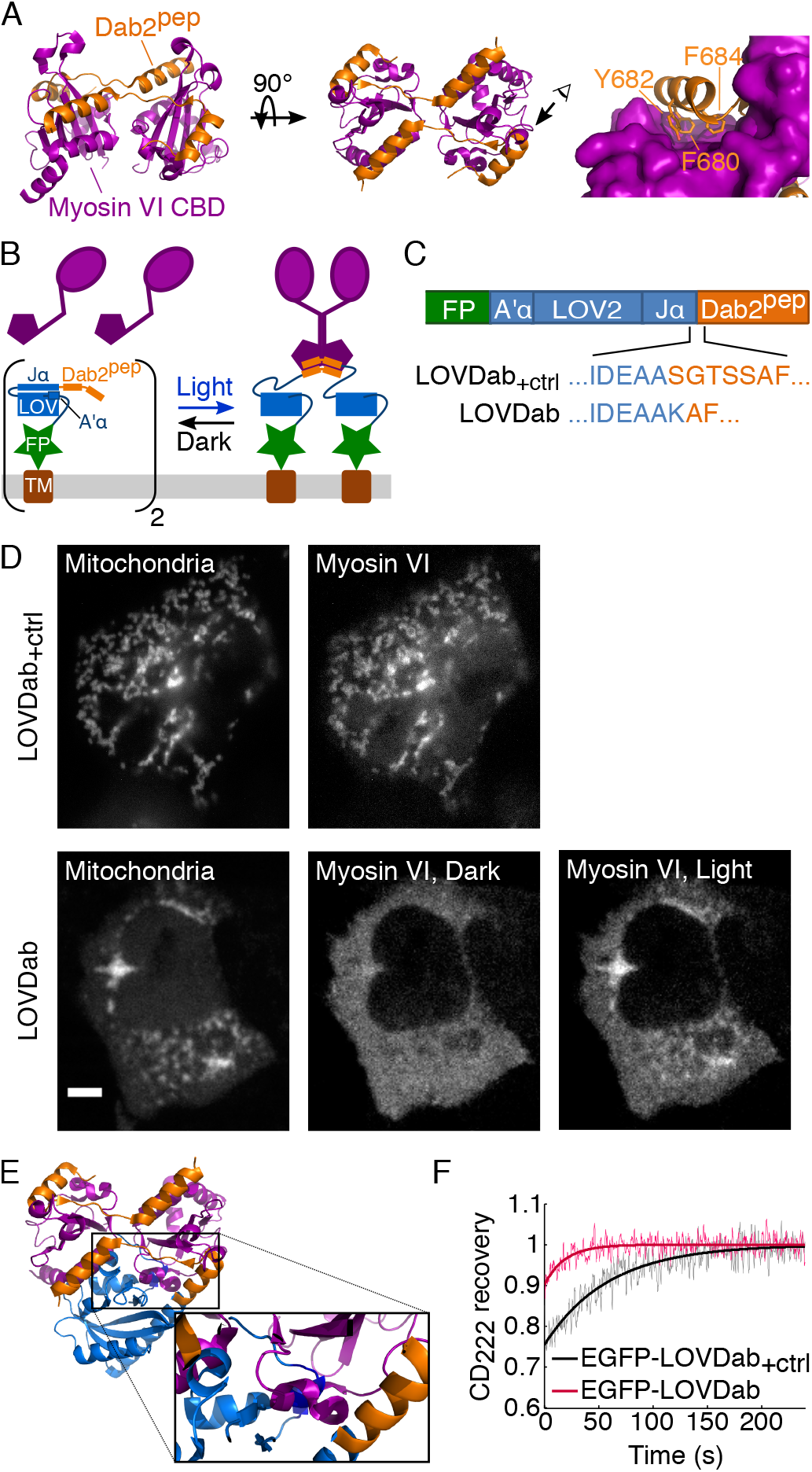
Design of LOVDab. (*A*) *Left and center*. The myosin VI CBD co-crystalized with Dab2^pep^ in two orientations illustrating how two Dab2^pep^ molecules straddle the CBDs. *Right*: View of N-terminal Dab2^pep^:CBD binding interface with aromatic residues shown in sticks. (*B*) Diagram of desired function of LOVDab. In the dark, Dab2^pep^ is prevented from binding myosin VI. When excited by blue light, the A’α and Ja helices of LOV2 unfold, releasing Dab2^pep^ to recruit myosin VI to the membrane. An N-terminal transmembrane (TM) tag targets LOVDab to a specific organelle in the cell. A fluorescent protein (FP) is used for visualization (*C*) Domain architecture of EGFP-LOVDab with the sequences of LOVDab_+ctrl_ and LOVDab at the fusion site. LOVDab_+ctrl_ is designed to constitutively activate myosin VI and is useful for optimizing LOVDab assays. (*D*) Recruitment assay from *B* performed using the constructs in *C*. LOVDab_+ctrl_ does not cage Dab2^pep^ in the dark whereas LOVDab only recruits myosin VI to the mitochondria when excited with blue light. Scale bar: 5 µm. (*E*) Alignment of LOV2 with Dab2^pep^:CBD crystal structure (with zoom to boxed region on right) suggests LOVDab obstructs myosin VI binding to Dab2^pep^ in the dark. Dark blue: A′α helix. (*F*) Time course of dark state photorecovery of constructs in *B* measured by circular dichroism (CD) at 222 nm normalized to their respective values in the dark state. At time = 0 sec the LOV2 domains are fully excited and allowed to recover in the dark.

We first tested whether Dab2^pep^ is sufficient to recruit full-length myosin VI using an uncaged, constitutively open fusion of Dab2^pep^ to the C-terminus of the LOV2 Jα helix (Fig. 1*C*). Residues 674-679 of Dab2^pep^ are mostly glycine and serine, suggesting that these residues could serve as a flexible linker between the LOV2 domain and the remaining residues in Dab2^pep^. This flexibility should prevent caging of the myosin VI binding site on Dab2^pep^ even when the Jα helix is folded in the dark. As expected, LOV2 fused to the full Dab2^pep^ sequence recruits myosin VI to mitochondria in the dark (Fig. 1*D*). Myosin VI is seen on puncta that correspond to the mitochondria visualized in the LOVDab_+ctrl_ channel. We refer to this construct as LOVDab_+ctrl_ (Fig. 1*C*), since it is always myosin VI binding-competent.

Having established that Dab2^pep^ is sufficient for myosin VI recruitment, we next sought to re-engineer LOVDab_+ctrl_ to have light-dependent binding. Our strategy focuses on caging the aromatic residues F680, Y683, and F684 on the Dab2^pep^’s N-terminal helix that are buried upon CBD binding (Fig. 1*A*, *right panel*). These three residues must be close enough to the Jα helix that they are sterically prevented from binding the CBD when the Jα helix is folded in the LOV2 dark state. This criterion is accomplished by reducing the number of intervening residues while maintaining a continuous helix encompassing the Jα helix and the N-terminal helix of Dab2^pep^. An additional variable is helical registry which controls the relative orientation of the Jα and the Dab2^pep^ helices, and therefore the angle of the CBD-binding residues in Dab2^pep^ with respect to the LOV2 domain. Additionally, disruption of the interaction between I539 on the Jα helix and the LOV2 core partially unfolds the Jα helix (37). These constraints together limit the amount of Jα sequence we can modify.

We tested a small battery of constructs for light-dependent myosin VI binding using our mitochondrial recruitment assay (Fig. 1*B*). One construct, which we refer to as LOVDab (Fig. 1*C*), exhibits robust recruitment of myosin VI in response to light (Fig. 1*D*). As intended, Tom70^helix^-FP-LOVDab localizes to the surface of mitochondria, which are visible as puncta (Fig. 1*D*). When the LOV2 is in the dark state, mCherry-myosin VI fluorescence shows a diffuse, cytosolic signature indicating that it is unbound and diffuse throughout the cytosol. Upon LOV2 activation via whole cell illumination with a blue, 488 nm laser, the myosin VI fluorescence is depleted from the cytosol and forms puncta corresponding to the mitochondria where Tom70^helix^-FP-LOVDab resides (Fig. 1*D*). Thus, LOVDab exhibits light-dependent binding to myosin VI.

Small deviations from this sequence abolish this light-dependent recruitment. LOVDab lacking K544 shows minor switching that was often accompanied with high dark state binding. LOVDab lacking both A543 and K544 shows weak, constitutive binding, indicating that moving the Dab2^pep^ closer to the LOV2 domain beyond these residues destabilized the Jα helix. The insertion of residues either C-terminal to K545 in the LOV2 sequence, or N-terminal to S678 in the LOVDab sequence, even when making compensatory insertions or deletions in the opposite fusion protein to maintain the myosin VI CBD binding site on the identical face of the Jα helix, prevents caging of Dab2^pep^. Adding S678 of Dab2^pep^ to LOVDab produced a construct with similar switching as LOVDab. Following the predicted register of the helix, this fusion would position F680, a residue that is buried in the Dab2^pep^:CBD interface (Fig. 1 *A*), facing toward the Hβ-Iβ loop of LOV2. However, this construct shows a small level of dark state binding, so we continued our study with LOVDab.

Both LOVDab and LOVDab_+ctrl_ contain the double mutation T406A/T407A on the A′α helix. This double mutation stabilizes the Jα helix by increasing the helicity of the neighboring A′α helix, which in turn improves caging of peptides fused to the Jα helix (26, 32). Modeling using the crystal structure of LOV2 and that of the Dab2^pep^_2_:CBD_2_ complex suggests that LOVDab blocks Dab2^pep^ from binding to the CBD of myosin VI by sterically occluding the CBD from its binding site on Dab2^pep^ (Fig. 1*E*). Interestingly, the largest steric clash is between the CBD and the A′α helix, suggesting that both terminal helices contribute directly to the caging of Dab2^pep^. In contrast, the flexible Dab2^pep^ residues 674-679 in LOV2Dab^pep^_+ctrl_ make this steric clash unlikely. This clash, together with the placement of Dab2^pep^ F680 in the LOV2 L546 position in LOVDab (38), may explain why this construct shows such robust light-dependent recruitment.

We further probed the structural changes in LOVDab and LOVDab_+ctrl_ using circular dichroism (CD) at 222 nm to interrogate the light triggered unfolding of the Jα helix and the uncaging of Dab2^pep^. In order to better replicate the protein used in the *in vivo* assays, we fused an EGFP to the N-termini of LOVDab and LOVDab_+ctrl_ for our CD measurements. The observed 25% fractional change in helicity in EGFP-LOVDab_+ctrl_ compares well with our previously measured values for the LOV2 domain (32), suggesting that fusion to either EGFP or Dab2^pep^ does not alter the extent of conformational change. We expect that Dab2^pep^ forms a helical extension of the Jα helix in LOVDab, whereas Dab2^pep^ is likely unfolded in EGFP-LOVDab_+ctrl_, as Dab2^pep^’s intrinsic helicity is only 0.64% (39). Consistent with this finding, the fractional change in helicity is smaller in EGFP-LOVDab than in EGFP-LOVDab_+ctrl_ (Fig. 1*F*), in part driven by the 20% higher intrinsic helicity of EGFP-LOVDab over EGFP-LOVDab_+ctrl_. This decrease in apparent unfolding of the Jα helix supports the notion that the Dab2^pep^ is more tightly caged in LOVDab than in LOVDab_+ctrl_.

### LOVDab controls myosin VI recruitment with high spatial and temporal control

One benefit of optogenetic approaches is their ability to control proteins at a subcellular level. Using a focused laser beam, myosin VI recruitment via LOVDab was performed selectively on subcellular regions (<5 µm in diameter) in a reversible and repeatable manner (Fig. 2*A–C*, Movie S1). Overlaying the helix recovery CD trace of LOVDab on the mitochondrial fluorescence (Fig. 2*C*) shows that myosin VI unbinds from the mitochondria at a similar rate to the Jα helix refolding in the dark, indicating that myosin VI recruitment is occurring through our LOV2 construct.

**Fig. 2.**
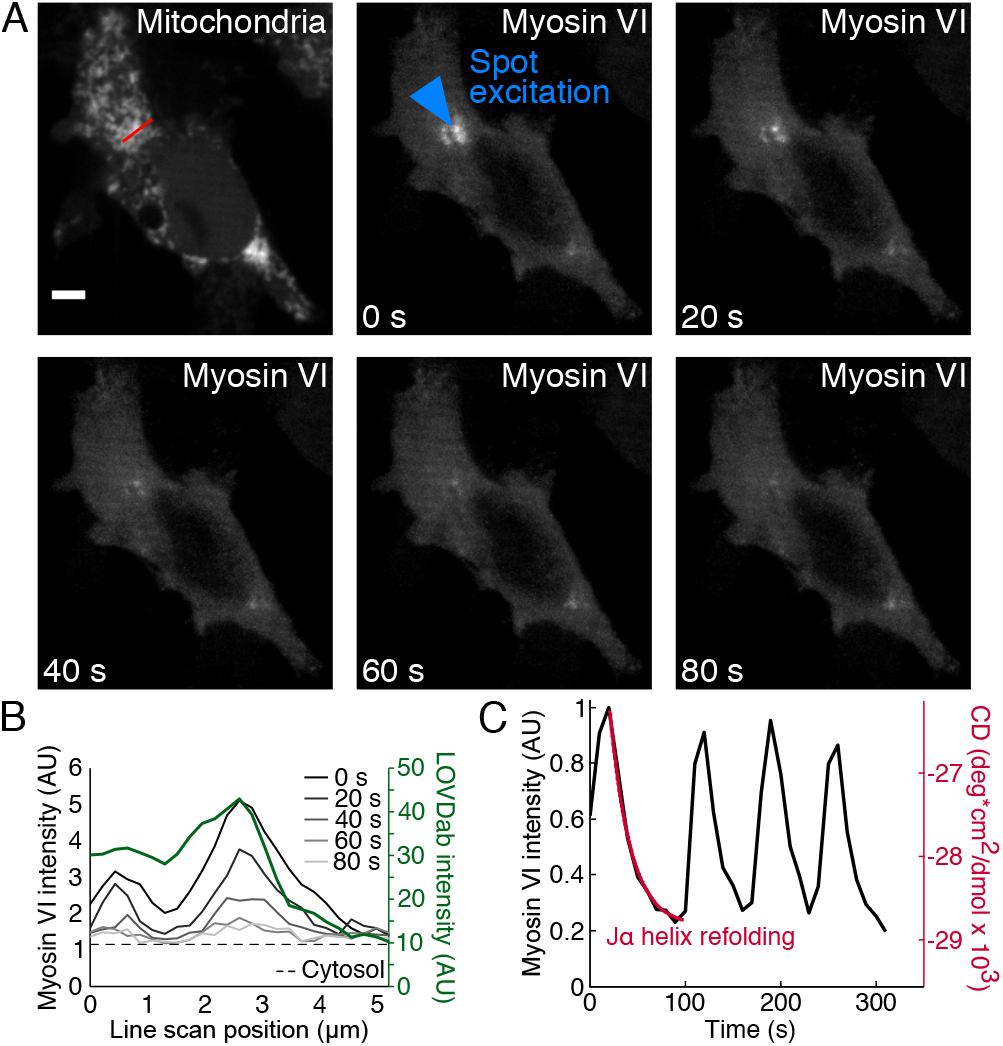
LOVDab recruits myosin VI to mitochondria with high spatial and temporal control. (*A–C*) Spot-specific, light-dependent recruitment of myosin VI to mitochondria. (*A*) Images of cell showing spot-dependent (blue arrow) recruitment of myosin VI. As cell is left in dark, myosin VI unbinds. Scale bar: 5 µm (*B*) Fluorescence intensity line scan across red line in *A* over time. (*C*) Mean myosin VI fluorescence in spot peak in *A* quantified over sequential pulses of light shows myosin VI recruitment is reversible and repeatable. LOVDab at spot are excited with blue light just before myosin VI images taken at t = 0, 10, 110, 180, and 250 sec in the plot. Myosin VI subsequently binds LOVDab at the spot (peaks), and unbinds as the cell is left in the dark (troughs). Overlayed circular dichroism (CD) trace of EGFP-LOVDab photorecovery indicates that the myosin unbinding rate is similar to the refolding rate of the Jα helix in LOVDab.

### Light-dependent recruitment of myosin VI stalls peroxisomes

Peroxisome movement is highly coupled to microtubule dynamics (40). Myosin V has previously been shown to slow the redistribution of peroxisomes upon recruitment of constitutively active kinesin-2, causing the peroxisomes to accumulate near the cell boundary. These observations suggest that myosins are capable of stalling and/or decoupling peroxisomes from microtubule machinery (41). We reasoned that if recruiting myosin VI to peroxisomes alters their motion inside the cell, this assay could determine whether LOVDab can activate myosin VI upon its recruitment. We targeted LOVDab and LOVDab_+ctrl_ to peroxisomes by fusing the Pex3 membrane targeting domain (Pex3^MTD^) to its N-terminus (42). We find that among cells co-transfected with Pex3^MTD^-FP-LOV2Dab2^pep^_+ctrl_ and myosin VI, cells having high myosin VI concentration exhibit constitutive binding between LOV2Dab2^pep^_+ctrl_ and myosin VI. In contrast, wild type peroxisomes do not show significant binding of myosin VI (Fig. 3*A* and *B*). Significantly, peroxisomes that bind myosin VI have overall slower velocities than wild type peroxisomes, supporting previous data showing that myosin V is capable of stalling microtubule-directed transport of these organelles (41).

**Fig. 3.**
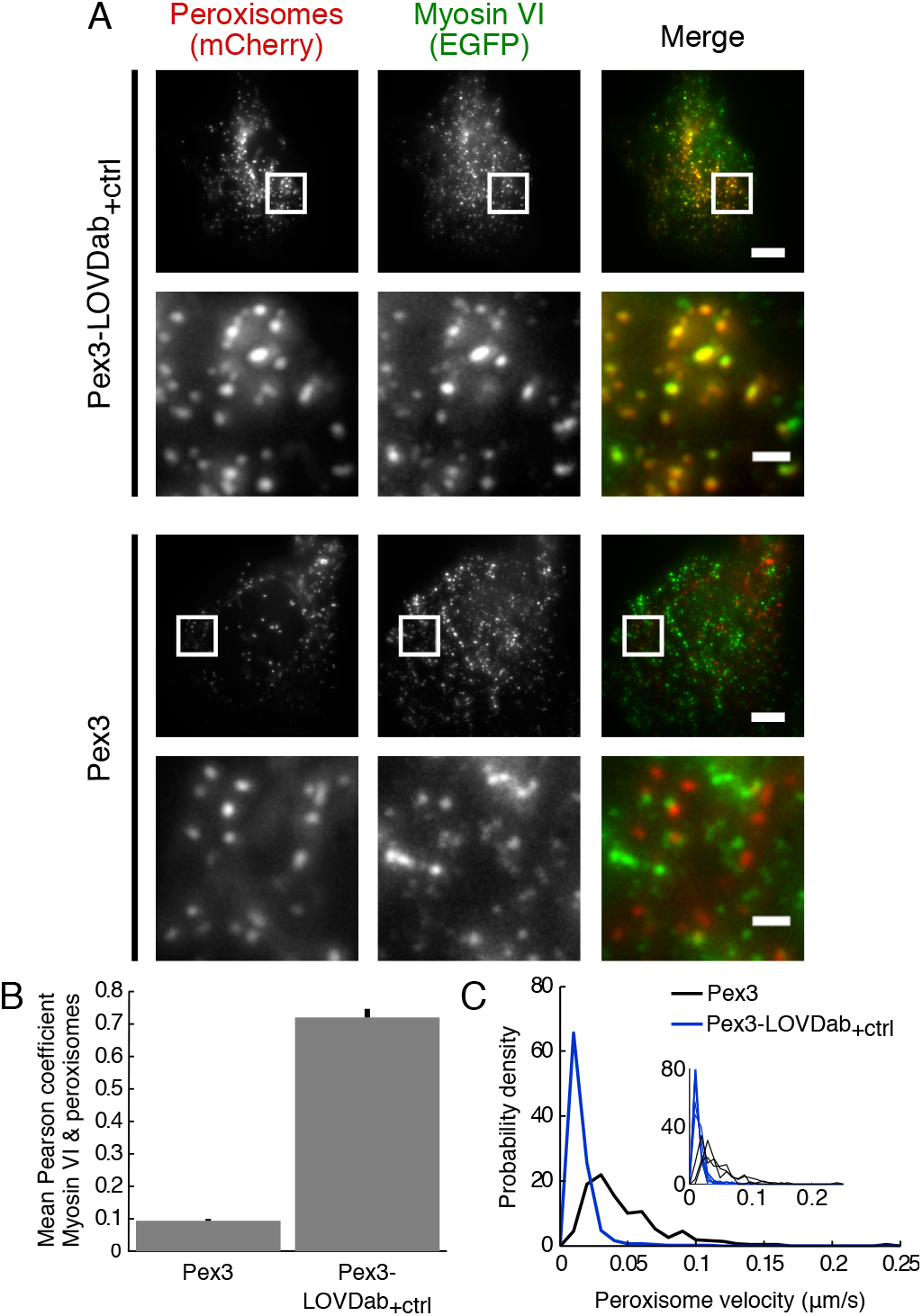
Myosin VI is recruited to peroxisomes via LOVDab. (*A*) The membrane targeting domain of Pex3 (Pex3^MTD^) is used to express either mCherry-LOVDab_+ctrl_ or mCherry only (wild type). *Top two rows*: Cells with peroxisomes containing LOVDab_+ctrl_ bind EGFP-myosin VI. *Bottom two rows*: Wild type peroxisomes lacking LOVDab_+ctrl_ do not bind myosin VI. The 2^nd^ and 4^th^ rows are expanded views of boxed regions in the 1^st^ and 3^rd^ rows, respectively. Scale bars: 5 µm (1 µm expanded). (*B*) Quantification of overlap between peroxisome and myosin VI channels confirms significant recruitment of myosin VI to peroxisomes labeled with LOVDab_+ctrl_ (mean ± SEM, p << .01, t-test; wild type, n=3; LOVDab_+ctrl_, n=6). (*C*) Peroxisome frame-to-frame velocities are markedly reduced in cells expressing Pex3^MTD^-mCherry-LOVDab_+ctrl_. Inset shows distributions for each cell used in analysis. Main graph shows distributions pooling all peroxisome data from these cells.

We next targeted LOVDab to peroxisomes, and were able to achieve reversible, light-dependent recruitment of myosin VI to this organelle (Fig. 4*A* and *B*; Movie S2). Furthermore, the recruitment occurs only to peroxisomes coated in LOVDab. Lit peroxisomes move more slowly (Fig. 4*C*), presumably because the recruited myosin VIs act as a dynamic tension sensor that stalls and anchors peroxisomes to the actin network (43). Reduced peroxisome movement is evident from standard-deviation projections of pixel values, as the fast moving peroxisomes form blurred patterns that coalesce into compact puncta in the light (Fig. 4*D*, Movie S3). After three minutes in the dark, the peroxisome motility resumes, as indicated by the blurred patterns reappearing. Moreover, peroxisome trajectories are more compact in the light (Fig. 4*E*), with a reduced diffusion coefficient (Fig. S2). We therefore conclude that LOVDab is capable of reversibly activating myosin VI *in vivo*.

**Fig. 4.**
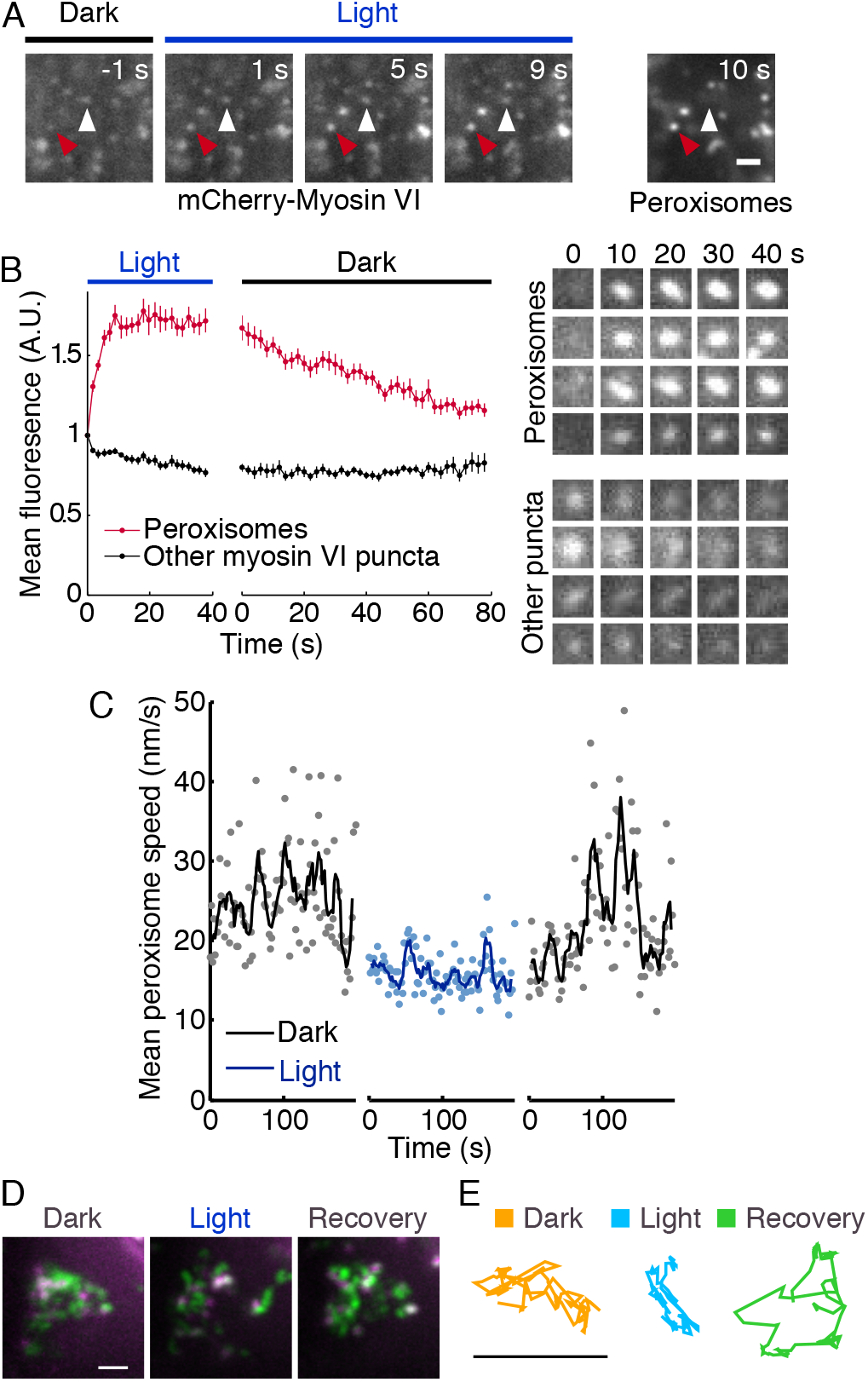
LOVDab reversibly recruits EGFP-myosin VI to peroxisomes in a light-dependent manner, which causes them to stall. (*A*) Images of a region of a cell showing light-dependent recruitment of myosin VI to peroxisomes. Vesicles showing light-dependent recruitment (red arrows) correspond to peroxisomes (*right image*) while vesicles binding myosin VI in the dark do not (white arrows). Thus, light-dependent recruitment occurs through LOVDab. Scale bar: 1 µm. (*B*) Quantification of mCherry-myosin VI fluorescence on vesicles in the cell pictured in (*A*). Robust, light-dependent recruitment of myosin VI occurs to LOVDab-labelled peroxisomes, and decays on other vesicles where LOVDab is not present. Individual vesicle traces were normalized to their dark state values just prior to LOV2 excitation (0 s, left panel) prior to calculating the mean (n=12, peroxisomes; n=7, other myosin VI puncta). Error bars are ±SEM. At right are images of four individual peroxisomes and four other vesicles over the excitation phase (image dimensions: 900 × 900 nm). (*C*) The mean frame-to-frame speed of peroxisomes decreases when the peroxisomes are illuminated. In total, 170-203 traces were collected per video. The line shows the smoothed data (5-timepoint rolling average). The mean speed is reduced in the illuminated period, and increases in the recovery period. These speed differences are statistically significant (p = 10^−81^ dark vs. light, and p = 10^−31^ light vs. recovery; Mann-Whitney U test over all unaveraged speeds). (*D*) Peroxisome excursions are reduced when illuminated. In green is a standard deviation projection of the peroxisome in the central region of Movie S4, showing the area covered by moving peroxisomes. The first frame of each period is shown in magenta, showing the starting position of the peroxisomes. Note the larger excursions in the dark and the recovery periods, vs. the light period (scale bar, 1 µm). (*E*) Representative trajectories of three peroxisomes, one from each period of the experiment. Note the lower extent of excursions in the light period. For an analysis of diffusion coefficients, see Figure S2.

### Activation of myosin VI in an *in vitro* gliding filament assay

Myosin VI could halt peroxisome movements by binding to actin filaments either with or without motor activity. In order to directly demonstrate activation of myosin VI motility by Dab2^pep^, we designed a modified gliding filament assay whereby we anchor an EGFP-LOV2-Dab2^pep^ fusion protein to the surface of a glass coverslip through an anti-GFP antibody (3E6). We then perfused soluble mCherry-myosin VI with actin together (Fig. 5*A*). This design allows us to interrogate myosin VI activation without altering its C-terminal cargo binding domain.

**Fig. 5.**
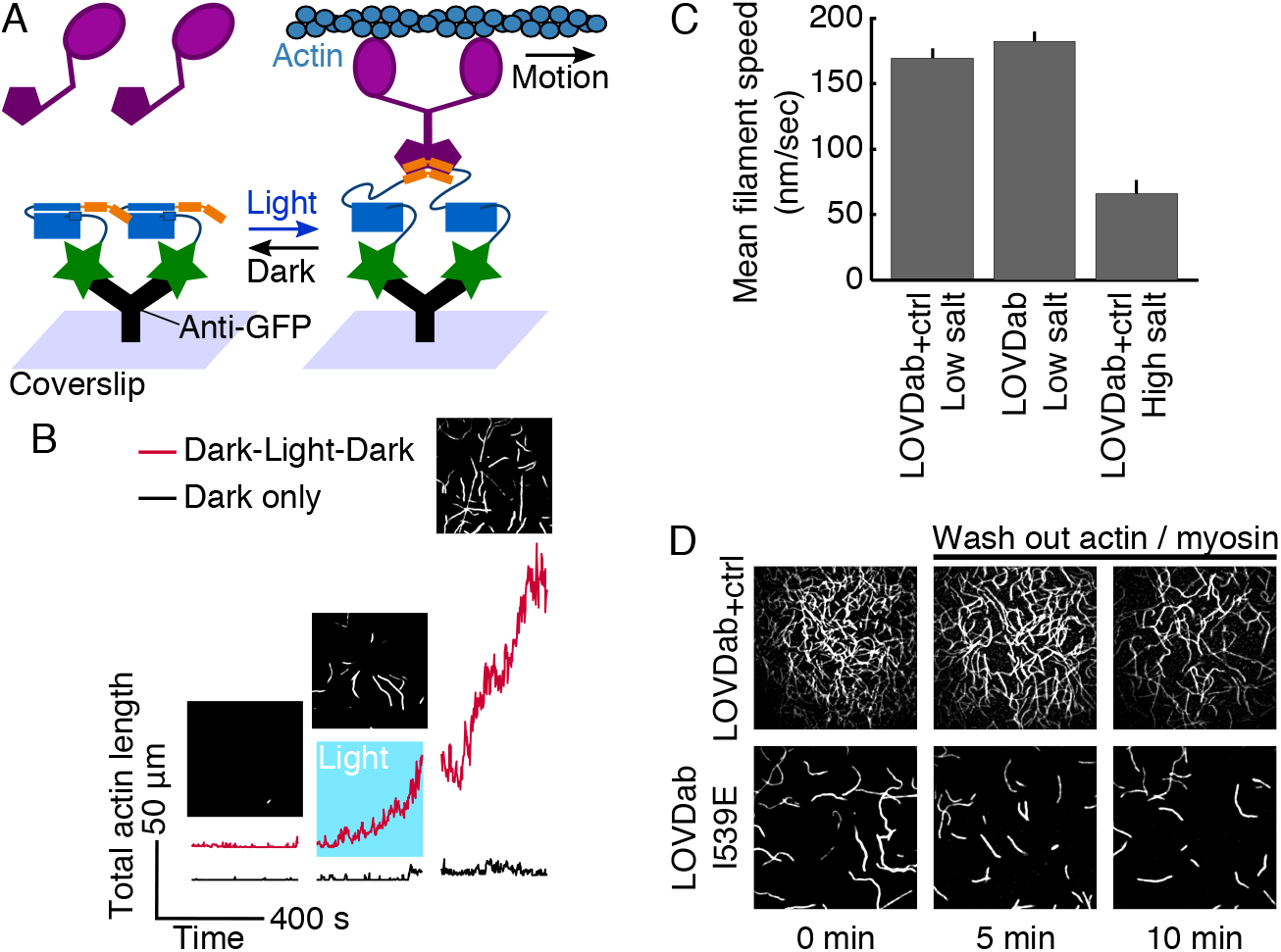
*In vitro* gliding filament assay indicates that LOVDab activates full-length human myosin VI in response to light. (*A*) Diagram of the modified gliding filament assay. EGFP-LOVDab is attached to the coverslip surface using an anti-GFP antibody. Soluble myosin VI is recruited to the surface by Dab2^pep^ and translates actin filaments. (*B*) Quantification of actin binding to surface over time for LOVDab. Two parallel slides were recorded, one with a photoexcitation period in the middle (red) and another that was not excited as a negative control (black). Images above the red trace are for the end of each period for the Dark-Light-Dark sample. (*C*) Filament speeds over various conditions. Salt significantly alters filament velocity while velocities for LOVDab and LOVDab_+ctrl_ are similar (mean + SEM; first column, n=4 videos; others, n=3). Data for LOVDab is pooled from videos recorded in light and returned to dark post-excitation (see methods). (*D*) *Upper row*: in high salt, actin releases from coverslip when free myosin and actin are washed out of the slide. *Lower row*: the I539E mutation unfolds the Jα helix, acting as a constitutive lit state mutant. This mutation allows LOVDab to recruit actin to the surface in high salt.

We find that full-length human myosin VI purified in a low salt ([KCl] = 25mM) MOPS buffer propels filaments at 170 nm/sec when recruited by LOVDab_+ctrl_. This velocity agrees well with published values (11, 44). Under these conditions, LOVDab+_ctrl_ recruits actin to the surface rapidly in the dark (Fig. 5*B*, *lower panels*; Movie S4).

When the coverslip is coated with LOVDab, myosin VI only recruits actin to the surface after the LOV2 is excited with blue light (Fig. 5*B*, Movie S5). After 400 seconds of excitation, the coverslip surface is not yet saturated with actin, indicating that the actin deposits at a slower rate than with LOVDab+_ctrl^−^_ coated coverslips. The velocity of the actin filaments recruited through LOVDab is similar to that of filaments recruited through LOVDab_+ctrl_ (Fig. 5*C*). This latter finding suggests that both constructs fully activate myosin VI. The slower deposition rate for LOVDab may be due to the lower availability of the Dab2^pep^ in LOVDab over LOVDab+_ctrl_, leading to fewer recruited myosin VI motors. It could also reflect the selective recruitment of microaggregates of myosin VI by LOVDab, which would diffuse to the surface more slowly than monomeric myosin VI (see below).

For both constructs, myosin VI appears to be recruited to the surface irreversibly, as once the LOV2 was activated, actin did not release from the slide (Fig. 5*B*). To determine whether this irreversibility was due to the high affinity of Dab2^pep^ for the myosin VI CBD, we attempted to invert the gliding filament assay to directly observe myosin VI walking on actin. However, we find that myosin VI purified in low salt solutions forms soluble microaggregates which are visible as diffusing clusters in the TIRF microscope. Testing a series of buffers, we find that myosin VI prepared in a high salt (KCl = 150 mM) Tris buffer does not form visible microaggregates. Myosin VI prepared in this second buffer is active in the gliding filament assay when LOVDab_+ctrl_ is on the surface. Consistent with a reduction in the myosin:actin affinity in high salt, filaments recruited to the surface in this assay have a significantly reduced velocity (Fig. 5*C*). This myosin also did not perform in the TIRF assay, either because the concentration needed to dimerize the myosin VI using cargo was prohibitively high for detection using TIRF, or because of the well-documented decrease in affinity between myosins and actin in high salt conditions (45, 46).

Unlike in low salt conditions, washing out free actin and myosin VI from the slide results in the slow release of myosin VI from LOVDab_+ctrl_ in high salt. (Fig. 5*D*) Together, the high off-rate of myosin VI from the mitochondria and the reversibility in the high salt myosin VI preparation suggest that the irreversibility seen in the gliding filament assays using myosin VI prepared in low salt was primarily due to the presence of myosin VI microaggregates. However, without the high avidity of the myosin VI microaggregates, we do not observe recruitment of actin to the surface via LOVDab under high salt conditions. We postulate that the lack of myosin VI recruitment to LOVDab *in vitro* is due to the engineered caging effect of the LOV2 domain on Dab2^pep^ in the LOVDab construct. This effect could weaken the effective K_D_ for binding of Dab2^pep^ to myosin VI to a point where even in the light, the myosin VI cannot be recruited at concentrations where it is soluble in our high salt buffers. Alternatively, the lifetime of the LOVDab:myosin VI:actin complex may be too short to propel the actin filaments. Importantly, LOVDab containing an I539E point mutation that unfolds the Jα helix (37) constitutively recruited actin in the presence of high salt myosin VI, proving that LOVDab is inherently capable of activating myosin VI (Fig. 5*D*).

We also tested whether a LOV2 domain fused to the myosin VI-binding residues (412-520) of optineurin (OPTN) could activate myosin VI in our gliding filament assay. These residues are predicted to form a coiled coil (47, 48). Consistent with this, EGFP-LOV2-OPTN_(412-520)_ migrates as a dimer in size exclusion chromatography (Fig. S1). However, it does not activate myosin VI. Lack of activation is consistent with recent studies showing that the large insert in the myosin VI tail domain, which is present in our construct, occludes the OPTN binding site (49). Furthermore, OPTN must be ubiquitinated prior to binding myosin VI (50).

### Multiple signals contribute to myosin VI activation

Other factors that may contribute to the differences in our *in vivo* and *in vitro* assays include the lack of Ca^2+^ and PI(4,5)P_2_ in the *in vitro* assays, both of which are known to bind myosin VI (18, 20). At myosin VI concentrations too low to interact with LOVDab_+ctrl_, the addition of PI(4,5)P_2_ rescues recruitment of actin to the coverslip (Fig. 6*A*). This rescue requires the presence of Ca^2+^ ions, suggesting that the three signals, PI(4,5)P_2_, Ca^2+^ and Dab2, are integrated to activate myosin VI in the proper time and place in the cell. Nevertheless, LOVDab is sufficient to recruit myosin VI and actin to the coverslip as shown in Fig. 5. We can therefore rule out mechanisms where myosin VI must first bind to PI(4,5)P_2_ and Ca^2+^ to expose an otherwise occluded Dab2 binding site.

**Fig. 6.**
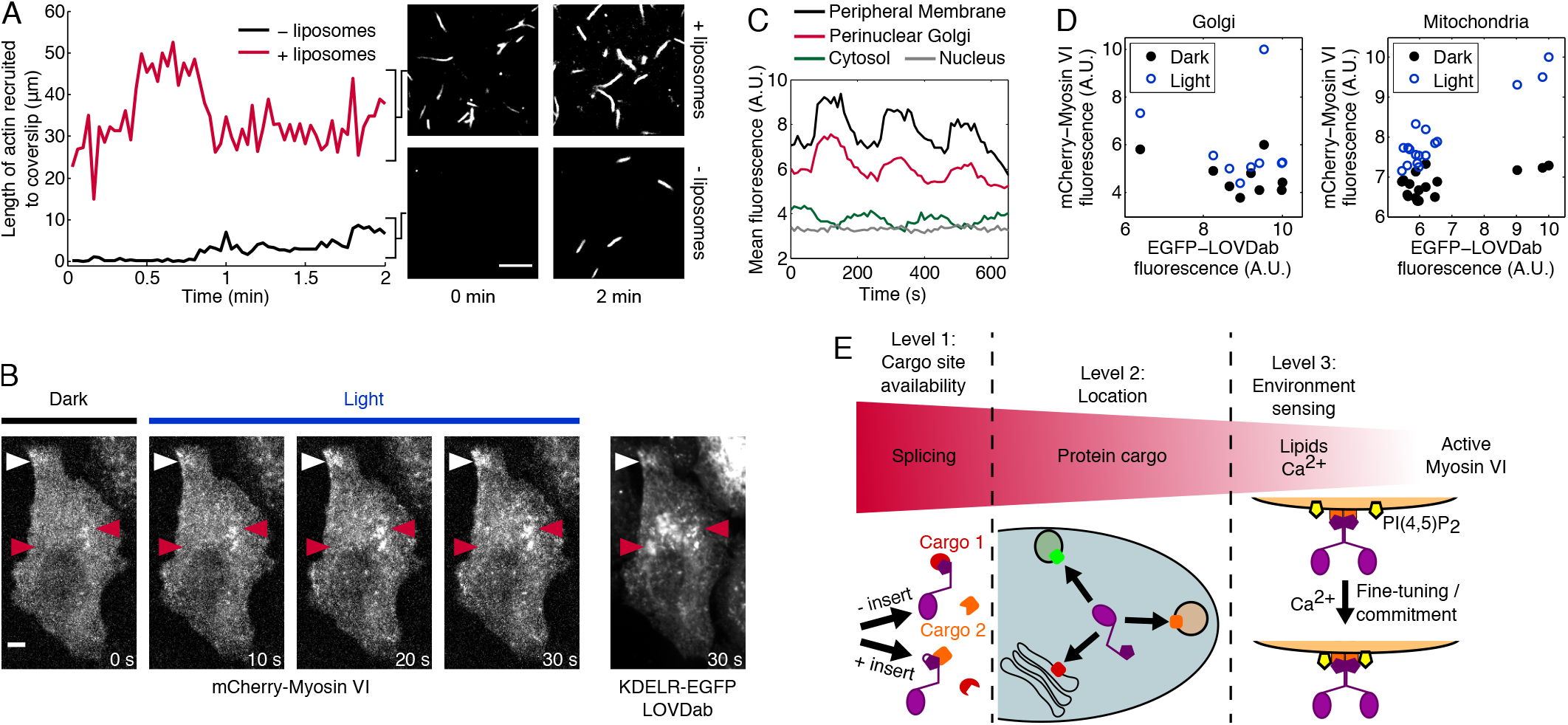
Myosin VI integrates cargo signals to deploy in site-specific manner. (*A*) At concentrations of mCherry-Myosin VI (prepared in physiological KCl conditions, see text) too low for LOVDab_+ctrl_ to recruit actin to the coverslip (- liposomes, *bottom*), addition of artificial liposomes containing 5% PI(4,5)P_2_ sensitizes full-length myosin VI to Dab2^pep^ cargo, rescuing recruitment of actin to the surface (+ liposomes, *top*). (*B*) LOVDab was targeted to the Golgi apparatus using a KDEL receptor (KDELR) with an R5Q/D193N double mutation to promote retention in the Golgi. White arrows highlight peripheral membrane that exceptionally recruits myosin VI. Red arrows highlight portions of perinuclear Golgi stacks, some of which recruit myosin VI well (right arrow) whereas others have minor switching (left arrow). Scale bar: 5 µm. (*C*) Mean fluorescence in different areas of the cell described in B. Perinuclear Golgi fluorescence is the mean of all perinuclear stacks. Cytosolic fluorescence is anticorrelated with Golgi fluorescence, indicating depletion of myosin VI from the cytosol upon excitation of LOVDab. Nuclear fluorescence is static and indicates the noise level of the measurement. (*D*) *Left*: The amount of myosin VI binding is not correlated with the amount of LOVDab on the different Golgi stacks in the cell shown in (*B*), suggesting other factors in the cell must alter the myosin VI:LOVDab affinity on the different membranes. *Right*: in a typical cell where LOVDab is targeted to mitochondria, the amount of myosin VI binding to mitochondria does correlate with the amount of LOVDab, suggesting that the additional factors influencing the myosin VI:LOVDab interaction on the Golgi are not present on mitochondria. (*E*) Model of myosin VI cargo integration in the cell. Alternative splicing biases myosin VI isoforms to different cargo proteins. Protein cargo recruits myosin VI isoforms to specific locations in the cell and determines its oligomeric state. Lipid, Ca^2+^, and other cargoes help myosin VI sense the environment, modulating its affinity for cargo, as seen in *A–D*.

This signal integration could explain recruitment of myosin VI to the Golgi via LOVDab with the KDEL receptor (KDELR). Overexpression of KDELR seems to be cytotoxic. At low expression levels KDELR-FP-LOVDab recruits myosin VI to the Golgi apparatus in a light-dependent manner (Fig. 6*B* and *C*; Movie S6). However, the amount of myosin VI binding to each LOVDab-labeled membrane does not correlate with the amount of LOVDab on that membrane (Fig. 6*B–D*). This lack of correlation cannot be explained by limiting myosin VI since when light is applied to the cell, additional myosin VI binds to each membrane. Thus, other factors on these membranes must alter the myosin VI affinity for its protein cargo Dab2^pep^, even between similar membranes. Plotting the same data for a typical cell showing light-dependent recruitment of myosin VI to mitochondria, which largely lack myosin VI PI(4,5)P_2_ (51), shows a different relationship. LOVDab and myosin VI levels on these membranes are correlated (Fig. 6*D*, *right*, Fig. S3), suggesting a linear dependence of myosin VI binding on the amount of LOVDab present. Together, these data suggest that myosin VI integrates multiple cargo signals to deploy in a site-specific manner in the cell (Fig. 6*E*, see discussion below).

## Discussion

Myosin VI is the only myosin that walks toward the pointed end of actin filaments (4, 5). This property gives it many unique roles throughout the cell including endocytosis, cell migration, and autophagy (52). Furthermore, the mechanism for myosin VI activation is still highly debated (12). To create a tool that could help address these questions, we engineered an optogenetic activator of myosin VI, LOVDab. Previous optogenetic efforts to control motors used two approaches. One used a clever optogenetic switch within the motor protein itself to achieve photoswitchable gearshifting (53). The other recruited constitutively active, truncated, forced-dimer motor proteins via engineered adapters (54, 55). These designs provide an excellent means of laser-induced force-generation *in vitro* and *in vivo*, and allow researchers to use members of each of the three classes of motor proteins. Our LOVDab differs in that it controls myosin VI through its native activation mechanism without any alterations to the myosin itself. Thus, our approach allows for direct tests of myosin VI function and control in cellular contexts. Additionally, since full-length myosin VI is auto-inhibited when not bound to cargo, our method couples recruitment and motor activity, preventing potential adverse effects on cytoskeletal systems when exogenous myosin VI is expressed *in vivo*.

Curiously, LOVDab places the residue F680 of Dab2^pep^ (res. 674-711) in the L546 position of LOV2, which Guntas *et al* observed to bind to a hydrophobic pocket in the non-FMN binding face of the β sheet of LOV2 (38). When optimizing their LOV2-based recruitment system, “improved Light Inducible Dimer (iLID)”, they noticed their optimal switch had a key phenylalanine residue that bound this pocket. Another parallel between our designs is a shortened photorecovery time. Their optimal construct exhibits a photorecovery time with τ ≈ 22 seconds relative to the wild type value of 80 seconds (32). Similarly, LOVDab exhibits a τ = 15 seconds compared to LOVDab_+ctrl_ which has a mean τ = 47 seconds, much closer to the recorded value for LOV2 406A/407A of 56 seconds (33). The relative difference in photorecovery times are evident from the rates of Jα helix reversion in Fig. 1*D*. Given the relative intolerance of LOVDab to deviations of F680 from this position, these results suggest that this hydrophobic pocket may be of general importance for caging peptides on the C-terminus of LOV2. Furthermore, a quickened dark state reversion time may highlight a key role of the L546 pocket in the overall conformational dynamics of the LOV2 domain. In any case, it seems that placing the Dab2^pep^ F680 in this pocket contributed to robust, light-dependent recruitment of myosin VI by LOVDab.

We demonstrated that LOVDab is capable of recruiting myosin VI in a light-dependent manner to mitochondrial, Golgi and peroxisomal membranes. Strictly speaking, however, binding of myosin VI to Dab2^pep^ is not equivalent to activation. Activation requires the release of autoinhibition of full-length myosin VI, allowing it to interact with actin in either a transport (56) or anchoring role (16). In order to demonstrate activation of myosin VI *in vivo*, we showed that myosin VI recruited to peroxisomes via our engineered positive control construct LOVDab_+ctrl_ stalls these organelles. Similarly, upon activation with light, LOVDab reversibly recruits myosin VI at high concentrations in the cell, stalling peroxisomes. These findings indicate that upon recruitment by LOVDab, myosin VI is capable of interacting with actin strongly enough to decouple peroxisomal motion from the underlying microtubule-based processes that drive their native behavior (40). This anchoring strongly suggests that LOVDab is capable of activating myosin VI *in vivo*.

In order to verify activation of myosin VI directly, we developed a modified *in vitro* gliding filament assay in which myosin VI is recruited to the coverslip via LOVDab or LOVDab_+ctrl_. Myosin VI purified under either high or low salt conditions is active when recruited through LOVDab_+ctrl_. Recent studies suggest that PI(4,5)P_2_ and Ca^2+^ transients may play roles in regulating myosin VI (18, 20). However, whether this regulation happens in conjunction with, or parallel to, activation by protein cargoes remains unclear. Our results suggest that Dab2-based activation does not require other signals, but that lipids can contribute as an independent binding partner. Whether there are additional inhibitory signals that block Dab2-based activation is currently unknown.

We recognized the ability of our *in vitro* assay with LOVDab to interrogate the interplay between these different cargo signals. We find that at low concentrations of myosin VI, PI(4,5)P_2_-containing liposomes enhance myosin VI’s affinity for LOVDab_+ctrl_ (Fig. 6*A*). This enhancement requires Ca^2+^. This requirement suggests that on membranes enriched in PI(4,5)P_2_ (57), Ca^2+^ transients act to enhance myosin VI’s effective affinity to its cargo protein Dab2 in the cell. Moreover, when LOVDab is targeted to the Golgi apparatus, the amount of myosin VI recruited does not scale with the amount of LOVDab on a given stack. Therefore, local differences must modulate myosin VI’s ability to bind these organelles. One cause for this difference could be in the concentrations of PI4P on the different stacks. Though myosin VI binds more strongly to PI(4,5)P_2_, it also shows significant binding to PI4P (18), which is found on the Golgi (57). Calcium transients may also regulate myosin VI recruitment to the Golgi, as secretory pathway organelles such as the Golgi are rich in Ca^2+^ (60). The cisternae in the Golgi have different luminal Ca^2+^ concentrations based on their developmental stage and can differentially release Ca^2+^. Thus, PI4P and Ca^2+^ are two putative signals that might control levels of myosin VI on the Golgi, independent of our LOVDab targeting system.

Although myosin VI recruitment levels do not correlate with the LOVDab quantity on the Golgi, they do correlate with the LOVDab quantity on mitochondria (Fig. 6*D*, *right*; Fig. S3). The signaling lipid PI(4,5)P_2_ has previously been detected on mitochondria (58), although at a much lower concentration than on the plasma membrane and in a sequestered form (58, 59). We are not aware of any report showing PI4P on mitochondria. Thus, we speculate that the differences seen in the LOVDab:Myosin VI binding patterns on the Golgi versus mitochondria could be partially explained by lipids and calcium modulating myosin VI’s affinity for the target.

Our findings are consistent with a model of myosin VI activation where, rather than having a given cargo for a given function, myosin VI integrates multiple signals at each membrane to modulate binding and motor activation (Fig. 6*E*). Regulation through splicing biases myosin VI to certain protein cargo adaptors (49). The protein cargos regulate the localization of myosin VI in the cell, and may also be the primary determinant of myosin VI’s oligomeric state (13). Once on-site, secondary cargo signals such as lipids or Ca^2+^ may fine-tune myosin VI’s activity at that location by increasing myosin VI’s effective affinity for its substrate, driving local activation (20).

LOVDab is a unique optogenetic tool in that it activates full-length myosin VI without altering the myosin itself, allowing for direct inferences on myosin VI activity and regulation *in vitro* and *in vivo*. Our approach here could guide similar optogenetic efforts to control native systems with minimal perturbation. LOVDab exhibits robust, light-dependent myosin VI recruitment to many different organelles, and myosin binding to LOVDab propels actin filaments *in vitro*. Using LOVDab, we discovered evidence for a new activation model for human myosin VI, whereby myosin VI integrates its activation signals to obtain a site-specific mode of activation. How each of these cargos are integrated, and how this integration is reflected structurally in myosin VI, are outstanding questions. In particular, the relation between the oligomeric state of myosin VI and its site-specific roles in the cell, and how this is impacted by different cargos, is not entirely known. We believe that LOVDab will be an extremely useful tool for answering these and additional questions about how this important motor protein functions in the cell.

## Materials and methods

### Plasmid construction

For protein purification, plasmids encoding LOV2-Dab2^pep^ fusion proteins were engineered within a pHISparallel1 vector encoding a modular -/NdeI/-His6-Protein G-/AgeI/-EGFP-/EcoRI/-LOV2-Dab2^pep^-/XhoI/- that allowed for facile swapping of domains via ligation. Full-length myosin VI was incorporated into baculovirus for expression in SF9 cells using the BestBac system by Expression Systems (Nevada, USA) according to the manufacturer’s guidelines. An mCherry2B (62) with an N-terminal FLAG tag was first incorporated at the myosin VI N-terminus within the pET vector (see below) prior to insertion into a modified pBlueBac plasmid via SLiCE cloning due to the large sizes of the fragments (63). We fused mCherry2B to its to improve solubility, reduce dimerization, and increase fluorescence of the protein over mCherry (62).

For live cell imaging, LOV2-Dab2^pep^ plasmid constructs were created for transfection using ligations from a modular pEGFP vector containing the general sequence -/NheI/-membrane tag-/AgeI/-FP-/EcoRI/-LOV2-Dab2^pep^-/BamHI/-Membrane tags include: for mitochondria, the mitochondrial outer membrane targeting sequence of yeast Tom70^helix^ (res. 1-40); for the Golgi Apparatus, full-length human KDEL receptor containing a R5Q/D193N double mutation to improve retention in the Golgi (64, 65); and for peroxisomes, the N-terminal targeting sequence of human Pex3 (res. 1-42) (Pex3^MTD^) with a (GGS)x3 linker (42, 66). Tom70^helix^ and B2AR were generous gifts from M. Glotzer. KDELR and Pex3^MTD^ were constructed from gBlocks ordered from Integrated DNA Technologies (Coralville, IA). Full-length human myosin VI, isoform 1 (a generous gift from M. Zhang) was expressed in their modified pET vector containing a sequence encoding -/NheI/-FP-/XhoI/-myosin VI-/NotI/-. Fluorescent proteins were exchanged via ligation.

### Protein purification

LOV2-containing constructs were purified as described previously (32). Briefly, constructs were expressed in E. coli with an (His)_6_-Protein G fusion to their N-terminus, followed by a TEV protease restriction site. After elution from the Ni-NTA column, proteins were dialyzed against a TEV protease cleavage buffer (50 mM Tris, 1 mM EDTA, 1 mM DTT, pH = 8.0) before or during the cleavage reaction. EDTA and DTT were removed by dialysis prior to passing the cleaved protein through a regenerated Ni-NTA column. Proteins were then concentrated and ran on a size exclusion column (HiPrep 16/60 S-100 HR, GE Healthcare Life Sciences) that was equilibrated in the relevant assay buffer.

Full-length human myosin VI with an N-terminal FLAG-mCherry2B fusion was incorporated into baculovirus as discussed above. SF9 cells were infected at a 1:1 MOI and proteins were harvested from SF9 cells 58-72 hours post infection. The cells were re-suspended in Lysis buffer (see below for recipes) containing 2 mM ATP, and a cocktail of protease inhibitors including 1mM phenylmethylsulfonyl fluoride (PMSF) and 10µg/mL each of: aprotinin, leupeptin, chymostatin, and pepstatin A. The cells were lysed by douncing and nutated for 40 minutes at 4°C to allow the myosin VI to diffuse out of the cells. The lysate was spun at 14,000 rpm in a Legend X1R centrifuge (F15S-8×50c rotor, Thermo Scientific) and a variable amount of anti-FLAG resin (M2, Sigma) added to the supernatant. After incubation, the resin was spun out of suspension at 900×g in a 5810R centrifuge (A-4-81 rotor, Eppendorf) and washed with wash buffer (see below) containing 3 mM ATP, a variable amount of Calmodulin, and protease inhibitors. Resin was repelleted using the same procedure and re-suspended in wash buffer before transferred to a drip column where the resin was further washed. Resin was then incubated with elution buffer (wash buffer + 0.2 mg/mL FLAG peptide) for 1 hour. Protein was eluted and dialyzed against wash buffer containing no ATP or protease inhibitors. In the low salt preps (see text) the lysis buffer was 50 mM Tris·HCl, 150 mM KCl, 0.5 mM DTT, and 0.1% Triton X-100, pH = 7.7. The wash buffer was 20 mM imidazole, 150 mM KCl, 5 mM MgCl_2_, 1 mM EDTA, and 1 mM EGTA, and 0.5 mM DTT, pH = 7.5. The storage/dialysis buffer was either AB: 25mM imidazole, 25 mM KCl, 4 mM Mg_2_Cl_2_, 1mM EGTA, 2mM DTT, pH = 7.5; or KMg25: 10 mM MOPS, 25 mM KCl, 1 mM EGTA, 1 mM MgCl_2_, and 1 mM DTT, pH = 7.0. For the high salt prep, the lysis buffer was 50 mM Tris·HCl, 150 mM KCl, 0.5 mM EDTA, 1 mM EGTA, 4 mM MgCl_2_, 0.5 mM DTT, and 0.1% Triton X-100, pH = 7.3, and the final dialysis/assay buffer was 50mM Tris·HCl, 150 mM KCl, 1 mM EDTA, 1mM EGTA, 5 mM MgCl2, and 0.5 mM DTT (1 mM DTT for final dialysis), pH = 7.3. For both preps, buffers were chilled to 0-4⍰C before use. pH values were set at room temperature.

Protein purity was assessed using SDS-PAGE. In the case of LOV2-containing constructs, protein quality was further assessed using UV-Vis spectroscopy to measure the FMN photorecovery rate and circular dichroism (CD) (see below) to measure the fractional change in helicity of the protein, as done previously in our lab (32). The UV-Vis spectroscopy was carried out on an Olis HP 8452 Diode Array Spectrophotometer (Bogart, GA). The recovery of FMN absorbance at 448 nm was fit to a single exponential using MATLAB to obtain the FMN dark state recovery time constant, τ_FMN_. Circular dichroism (CD) was performed on a Jasco 715 spectrophotometer and a 40W white LED lamp (BT DWNLT A, TheLEDLight.com) was used to excite the LOV2 domain in the cuvette for ≥ 12 seconds for photo excitation. The refolding traces at 222 nm were fit to a single exponential using MATLAB and the fractional change in helicity in the protein as δ_222_ = (CD_222,dark_-CD_222,light_)/CD_222,dark_.

### Gliding filament assay

Nitrocellulose-coated coverslips were first coated in anti-GFP antibody (clone 3E6 MP Biomedicals, California, USA) at 0.25 µg/mL in PBS. Following this, coverslips were blocked using 1 mg/mL BSA (ELISA grade, EMD Millipore) and the antibody saturated using EGFP-LOV2-Dab2^pep^ at 1 µM. After this, the slide was washed using KMg25 and a solution containing myosin VI, F-actin, and an oxygen scavenging system (0.45% glucose (w/v) 0.5% (v/v) BME, 432 µg/mL glucose oxidase, and 72 µg/mL catalase) (67) was perfused. Slides were then imaged using an ×100, 1.65 NA objective (Olympus) on a custom-built total internal reflection microscope employing an EMCCD camera (iXon; Andor Technologies). This microscope was controlled with the open source Micro-Manager program (www.micro-manager.org). For assays involving lipid cargo, artificial liposomes from Echelon Biosciences (Utah, USA) containing 5% of the signaling lipid PI(4,5)P_2_ were added to the assay buffer. The liposomes had no effect on myosin VI activity if Ca^2+^ was not present at a total concentration of at least 1.5 mM (corresponding to a free Ca^2+^ concentration of approximately 25 µM). The necessity of Ca^2+^ matches previous results but this concentration is approximately tenfold lower than previously used for assessing the interaction of lipids with the myosin VI tail domain (18).

Gliding filament velocities in Fig. 5*C* were calculated from at least 3 videos from at least 2 independent preparations of myosin VI, using at least 10 filaments/video and at least 10 frames/filament. MTrackJ was used to generate the actin tracks (68). Only filaments showing smooth and continuous movement were tracked, and only over frames where filament ends were in view. Myosin VI recruited by LOVDab in the light in low salt continued to propel filaments after returned to the dark. Since actin velocities did not differ largely between light and dark, these data were pooled to calculate the mean velocity in Fig. 5*C*. To quantify the length of actin on the coverslip, images were background subtracted and thresholded to create binary images of the actin that were then skeletonized using the ImageJ function. The sum of pixels in the images were converted to length using the pixel calibration and used as the estimate for the length of actin on the coverslip.

### Live cell assays

HeLa cells were passaged in Dulbecco’s modified Eagle medium (DMEM) from Corning (New York, USA) supplemented with 10% heat-inactivated FBS (Sigma, Missouri, USA) and 1x HyClone pen-strep-L-glut (GE Healthcare Life Sciences, Utah, USA). One day prior to transfection, cells were split onto plasma cleaned glass coverslips. Coverslips that were 80-95% confluent were washed into DMEM media lacking supplements 10-20 minutes prior to transfection. Cells were co-transfected with a combination of plasmids containing genes for either an FP-LOV2-Dab2^pep^ construct fused to a gene encoding an organelle-targeting transmembrane protein at its 5’ end or a gene encoding full-length human myosin VI with either EGFP or mCherry2B (see above) at its N-terminus using Lipofectamine 2000 (Life Technologies, Illinois, USA), according to the manufacturer’s guidelines. Cells were imaged 18-24 hours post-transfection. Confocal images were taken using a 63×, 1.4 numerical aperture (NA) objective on an Axiovert 200M microscope (Zeiss) with a spinning disk confocal (CSU10, Yokogawa) and an electron-multiplying charge-coupled device (EMCCD) camera (Cascade 512B, Photometrics). The microscope was controlled using MetaMorph (Molecular Devices). TIRF images (peroxisome experiments) were taken using the microscope used in the gliding filament assay. Videos were background subtracted using either the rolling ball or parabolic algorithms in ImageJ (69). Quantified pixel values are plotted post-background subtraction to match images shown.

For the peroxisome studies, particle tracking was performed using the TrackMate plugin for FIJI and analyzed in MATLAB, Julia, and R. More than 300 peroxisome traces from three wild type cells and >950 peroxisome traces from six Pex3™-FP-LOVDab_+ctrl_ cells were used for the data in Figure 3*C*. Each trace included at minimum ten frames. These cells were chosen based on an arbitrary fluorescence cutoff for myosin VI and LOVDab as peroxisomes in LOVDab_+ctrl^−^_-expressing cells with low levels of either protein behaved similarly to those in wildtype cells. As previously noted (40), peroxisomes in the interior of the cell are largely immobile, so these were excluded from analysis. The final data collected in Figure 4*C* and Figure S2 include 170-203 peroxisome traces per video.

## Author Contributions

ARF, TRS and RSR designed research; ARF performed research and analyzed data, ARF, TRS and RSR wrote the manuscript.

## Acknowledgments

The authors thank Michael Glotzer and Margaret Gardel for sharing equipment, and members of the Sosnick and Rock labs for helpful discussions. This work was supported by NIH R01s GM078450/S1 and GM109863 (to RSR), GM55694 (to TRS) and NIH T32 GM007183 (to B. Glick).

